# Pathogen-induced red pigmentation uncovers a conserved floral defense in Asteraceae

**DOI:** 10.64898/2026.02.24.707787

**Authors:** H.M. Suraj, Lisa-Marie Roijen, Malleshaiah SharathKumar, Justin J.J. van der Hooft, Boas Pucker, Jan A.L. van Kan

## Abstract

Flower colour is predominantly selected for visual appeal and pollinator-driven evolution, however its role in plant-pathogen interaction remains unclear. Here, we investigated the relationship between anthocyanin accumulation and resistance to the necrotrophic pathogen *Botrytis cinerea* in *Chrysanthemum morifolium*. Using a unique set of spontaneous colour mutants sharing a common genetic background, we demonstrate that higher anthocyanin levels are associated with enhanced resistance to *B. cinerea*. Furthermore, we also observed pathogen induced anthocyanin accumulation at pathogen penetration sites in the white cultivars. Transcriptomics and metabolomics analyses revealed shift in metabolic pathways from flavonols towards flavone and anthocyanin biosynthesis during infections. A broader survey of taxonomically distant flowering species identified pathogen-induced red pigmentation is largely restricted to the members of the Asteraceae family. Notably, we observed divergence between wild and domesticated species, where the wild species retained the inducible pigmentation response and exhibited resistant to *B. cinerea*, whereas the cultivated species showed loss of this response, correlating with its increased susceptibility. Overall, our findings indicate that breeding for ornamental traits may have inadvertently selected against floral immunity.

## Introduction

Flowering marks a critical transition in the plant life cycle, shifting developmental priorities toward reproduction. Despite their transient nature and rapid senescence post-anthesis, floral organs must remain protected from pathogen attack to ensure reproductive success. This process of physiological decline, characterized by nutrient remobilization and cell death, renders floral tissues significantly more susceptible to diseases compared to other vegetative organs. This knowledge gap is particularly relevant in horticultural systems, where disease incidence after harvest often further shortens the already limited post-harvest lifespan of cut flowers.

In ornamentals, flower colour is predominantly selected for consumer appeal. However, the role of floral pigments in disease resistance has received limited attention. Anthocyanins, the pigments responsible for red, pink, and purple coloration, are known to contribute to plant defense against both biotic and abiotic stresses (Grünig et al., 2025). This dual function is particularly relevant for the Chrysanthemum (*Chrysanthemum morifolium* cv. Ramat), the second most traded cut flower globally, after rose, and renowned for its extensive diversity in colour, shape and morphology of flowers (SharathKumar & Heuvelink, 2026). Despite its economic importance, it remains unclear whether variation in flower colour, specifically the concentration of anthocyanins influences the floral susceptibility to diseases.

To address this knowledge gap, we investigate how these pigments affect the susceptibility to necrotrophic plant pathogen *Botrytis cinerea*. This fungus is one of the most destructive plant pathogens worldwide, infecting more than 1,400 plant species and causing severe economic losses in the cut-flower industry (Elad et al., 2016). Disentangling the specific effects of floral colour from other genetic traits is challenging in heterogeneous diverse populations. Therefore, we utilized a set of spontaneous colour mutants derived from the commercial chrysanthemum cultivar ‘ResQ’. These lines share a common genetic background but differ substantially in petal pigmentation, including dark pink, salmon, light pink, pink, and white phenotypes. By quantifying anthocyanin concentrations and assessing the susceptibility of flowers following *B. cinerea* inoculations, we aimed to determine whether higher anthocyanin concentration is associated with reduced susceptibility.

Besides constitutive pigmentation differences, floral pigments can also be part of an inducible defense response. Recently, we reported that infection by *B. cinerea* can trigger localised anthocyanin and flavonoid accumulation at infection sites in the wild species *Chrysanthemum seticuspe*, a response directly linked to resistance (Suraj et al., 2025). However, it remains unknown whether cultivated species *Chrysanthemum morifolium* cv. Ramat (hybrid hexaploid) retains such ability to mount an infection-induced pigmentation response. To investigate this, we combined microscopy, transcriptomics, and metabolomics to examine infection-induced pigmentation in both coloured and white *C. morifolium* cultivars. Finally, we extended our analysis to several white-flowered species, including both cultivated and wild representatives across angiosperms, to test whether *B. cinerea* infection can overcome (epi)genetic blocks in the anthocyanin biosynthetic pathway, and restore pigment production in otherwise non-pigmented flowers.

## Materials and Methods

### Plant material

*Chrysanthemum morifolium* cuttings were obtained from Deliflor Chrysanten and Dekker Chrysanten (The Netherlands). Cuttings were treated with indole-3-butyric acid (IBA) rooting hormone and placed in a peat-based commercial potting substrate under high humidity for 2 weeks to induce rooting. Rooted cuttings were then transplanted into 1.5 litre pots filled with the same substrate and grown inside the glasshouse at Unifarm (Wageningen University & Research, Wageningen, The Netherlands) at 21°C (day), 19°C (night), and ∼60% relative humidity. Short-day conditions were imposed using blackout screens from 17:00 to 07:30 (14.5 h dark period; 9.5 h light period). Supplemental high-pressure sodium (HPS) lighting (Master GreenPower cgt 400 W, Philips) provided a canopy level Photosynthetic Photon flux density (PPFD) of approximately 200 µmol m⁻² s⁻¹ during the light period. Lamps were switched on when outside global radiation (GR) dropped below 150 W m⁻² and switched off when GR exceeded 250 W m⁻².

### *B. cinerea* culture and flower inoculations

*Botrytis cinerea* strain B05.10 was cultured on malt extract agar (MEA) at 20°C for 10 days under a 16 h light/8 h dark photoperiod. 20 mL of sterile Milli-Q (MQ) water was added to each plate, and conidia were scraped from the surface using a sterile spatula. The resulting mycelial and conidial suspension was filtered through glass wool and transferred to a 50 mL tube. The suspension was centrifuged for 10 min at 1000g to wash the conidia. The spore concentrations were determined by counting using a haemocytometer and later adjusted to 10⁷ spores mL⁻¹. Fully opened flowers were used for *B. cinerea* inoculations. The petals were inoculated with 2 µL drops containing 100 spores µL⁻¹. The inoculation media consisted of Gamborg B5 (Duchefa, The Netherlands), containing 3.2 g L⁻¹ minerals and vitamins (Gamborg et al., 1976), supplemented with 10 mM sucrose and 10 mM potassium phosphate (pH 6.0). For inoculations on petunia, sunflower and gerbera we used water instead of inoculation media because they were highly susceptible and tissue would otherwise be necrotic within 12-16 hours.

### Confocal microscopy

Leica DMI-8 system paired with a Stellaris-5 confocal laser scanning microscope was used to visualise the pigmentation response along with the fungal infection structure. Petals were inoculated with a GFP-tagged strain of *B. cinerea* B05.10 (Schumacher, 2012) and observed at 1 dpi. Using a white-light laser, samples were excited at 561 nm and emission spectra were recorded between 600-650 nm, which is the appropriate range for detection of anthocyanins without including chlorophyll auto-fluorescence signals previously described by (Kallam et al., 2017).

### Tissue staining

Staining of inoculated petals at 1 dpi with p-Dimethylaminocinnamaldehyde (DMACA) was performed as described by Li et al., 1996 with the omission of the decolouration step as the tested petals were already white. A freshly cut slice of apple was used as positive control. This staining allowed to visualise the potential presence of proanthocyanidins, also known as condensed tannins.

### qPCR analysis and primer sequences

Total RNA was extracted using the Maxwell® RSC Plant RNA Kit (Promega, USA), and cDNA was synthesized with MMLV reverse transcriptase (Promega, USA). qPCR was performed with the SensiFAST SYBR No-ROX kit (OriGene) on a Bio-Rad CFX96 Real-Time System (C1000 Touch). Primer sequences (5′–3′) were: Cs_actin_F CAGGATGAGCAAGGAAATCACC, Cs_actin_R AGGTGCTGAGTGATGCAAGGAT, Cs_ANS_F CGTCGGGGAAGATTCAAGGGT, and Cs_ANS_R AGCAGGTATGTAATCGCTAGGCG. Ct values were averaged per sample, normalized to Actin, and relative expression was calculated as 2^-ΔCt^.

### Anthocyanin quantification

Petal tissue was freeze-dried, ground to a fine powder, and an aliquot of dry material was weighed. Anthocyanins were extracted in acidified 75% methanol (methanol:water, 75:25, v/v) containing 1% HCl (v/v) by sonication for 20 min, followed by centrifugation at 13,000 rpm for 10 min. The supernatant was collected and absorbance was measured at 520 nm. Quantification was performed using an standard calibration curve prepared with cyanidin-3-O-glucoside (C3G) standard.

### Sample preparation and transcriptomics analysis

To ensure comparable developmental and growth conditions for transcriptomic sampling, cuttings were obtained from the respective breeding companies and grown under the same conditions. Flower tissues were collected in Wageningen (The Netherlands) in spring 2025. Samples of petal inoculated with *B. cinerea* or mock-inoculated were taken at different timepoints using 5 mm disposable biopsy punches (Robbins Instruments) to isolate a narrow disc of tissue around the site of inoculation. The collected discs were immediately frozen in liquid nitrogen and stored at -20°C. Three biological replicates were collected per timepoint, with each replicate consisting of petals from at least 3-4 flower heads. The collected samples were freeze-dried and ground to a fine powder in a Retsch MM 400 mixer mill for 10 minutes at 30 Hz using frozen Retsch blocks. For each biological replicate, 5 mg of powder was weighed for RNA isolations. Total RNA was extracted using the Maxwell 16 LEV Plant RNA Kit (Promega, USA). RNA-seq was performed with sequencing depth of 10 million reads per sample by the Beijing Genomics Institute (BGI, Shenzhen, China). All the raw data and metadata is publicly available (see data availability section). Clean reads were mapped to the *Chrysanthemum morifolium* cv. ‘Zhongshanzigui’ (yellow or slightly orange coloured flower petals) reference genome sequence (Song et al., 2023) using HISAT2 v2.2.1 with default parameters (Kim et al., 2019). Transcript quantifications were conducted with StringTie v3.0.3 (Pertea et al., 2015), and the read counts were normalized using DESeq2 v1.42.1 in R v4.3.3 (Love et al., 2014).

### Functional annotation of genes in *C. morifolium*

Flavonoid biosynthetic genes in *C. morifolium* were predicted using KIPEs v3.2.6 (Rempel et al., 2023) with default parameters. Candidate genes were selected based on the highest similarity scores, with priority given to the genes that contained all conserved amino acid residues and functional domains. When multiple candidates for a given enzyme did not share fully conserved residues, the top-scoring sequence was retained. Transcription factors were identified using bHLH_annotator v1.04 (Thoben & Pucker, 2023) and MYB_annotator v0.235 (Pucker, 2022) with default parameters.

### Sample preparation for metabolomics experiments

Metabolites from ResQ flowers were extracted by harvesting fully open inflorescences, freeze-drying the petals, and grinding them to a fine powder using a Retsch mill. For each sample, 5 mg of powder was extracted with 150 µL of 75% methanol / 25% dH₂O (v/v) containing 0.2% formic acid, followed by 20 min sonication and 10 min centrifugation at maximum speed. The supernatant was transferred to glass vials, sealed, and used for LC–MS/MS analysis. To extract the metabolites from the white flowers infected with *B. cinerea*, instead of grinding the tissues, we sampled the inoculation droplet from the infected petal surface as the compounds would leak from the tissue and diffuse into the droplet (2-dpi). The liquids from each biological replicates were pooled and extracted with a polar solvent consisting of 75% methanol and 25% dH₂O with 0.2% formic acid. Mock samples were prepared in the same way but without the fungal inoculations. A 100 μL volume was transferred to glass vials and sealed for mass spectrometry analysis. Around 22 flavonoids standards were run at the same time and are included in the metadata file of the LC-MS/MS analysis.

### LC-MS/MS data acquisition

LC-MS/MS was conducted using a Vanquish UHPLC with Exploris120 Orbitrap system (Thermo Scientific), comprising with an Vanquish photodiode array detector (220–600 nm) connected to an Orbitrap Exploris 120 mass spectrometer equipped with an electrospray ionization (ESI) source. The injection volume was 10 μl. Chromatographic separation was performed on a reversed-phase column (Acquity UPLC BEH C18, 1.7 μm, 2.1 × 150 mm; Waters) at 40°C. Degassed eluent A [ultra-pure water: formic acid (1000:1, v/v)] and eluent B [acetonitrile: formic acid (1000:1, v/v)] were used at a flow rate of 0.4 mL/min. For analysing anthocyanins, a steeper linear gradient of 5 to 35% acetonitrile (v/v) over 20 minutes was applied, followed by washing and equilibration. FTMS full scans (m/z 90.00–1350.00) were recorded at a resolution of 60,000 FWHM. maximum injection time of 100 ms. The ESI source parameters were set to a spray voltage of 2.5 kV (positive-ionization mode), and a capillary temperature of 290 °C. Data-dependent MS² spectra were acquired for the four most intense precursor ions per scan at a resolving power of 15,000 (m/Δm), using an AGC target of 1 × 10⁵, a maximum injection time of 118 ms, and an isolation window of 1.0 m/z. Fragmentation was achieved using stepped normalized collision energies (NCE) of 20, 40, and 100 eV (%) to generate diagnostic fragment ions supporting the annotation. All the datasets were obtained with positive ionization as we were mainly interested in flavonoids and anthocyanins and these compound classes are both ionizing well in positive mode.

### LC-MS/MS data analysis

Raw Thermo (.raw) files were converted to mzXML using MSConvert (ProteoWizard v3.0.24207; (Chambers et al., 2012). Feature detection was done in mzmine v4.4.3 (Schmid et al., 2023) using the wizard in positive ion mode. Data were cropped to 2.00–20.00 min, with chromatogram smoothing turned on. Mass detection used a factor-of-lowest-signal noise threshold (MS1 5.00, MS2 2.50) and a minimum feature height of 5E4. The ADAP chromatogram builder was used with minimum consecutive scans 5, minimum intensity for consecutive scans 1E4, minimum absolute height 5E4, max peaks per chromatogram 15, approximate feature FWHM 0.10 min, and scan-to-scan m/z tolerance 0.0030 m/z or 10 ppm. Chromatograms were smoothed using Savitzky–Golay. Features were resolved using the local minimum feature with chromatographic threshold 83.3%, minimum search range 0.10 min, minimum absolute height 5E4, minimum peak top/edge ratio 2.00, peak duration range 0.00–3.01 min, and minimum scans 5. Deisotoping was performed using the ¹³C isotope filter (m/z tolerance 0.0030 m/z or 10 ppm, RT tolerance 0.10 min, monotonic shape on, maximum charge 2, representative isotope = most intense; never remove features with MS2). Feature lists were aligned using the join aligner with sample-to-sample m/z tolerance 0.0030 m/z or 10 ppm (weight for m/z 3) and RT tolerance 0.20 min (weight for RT 1). The mzmine batch file is provided on Zenodo (see Data availability).

Features detected in mzmine were exported as an .mgf file and a feature quantification table and used for Feature-Based Molecular Networking (FBMN) in GNPS2 (gnps2.org) (Nothias et al., 2020; Wang et al., 2016)) using the FBMN workflow. The precursor ion mass tolerance was set to 2.0 Da and the fragment ion mass tolerance to 0.5 Da. The molecular network was constructed with a minimum cosine score of 0.7 and a minimum of 6 matched fragment ions between spectra, with a maximum precursor mass shift of 1999 Da. Network topology settings were: maximum molecular family size 100 and maximum number of neighbour nodes per feature (topK) 10. Feature abundances were normalized by row sum (RowSum). Spectral library matching used a minimum cosine score of 0.7 with at least 6 matched peaks; analog searching was off. The FBMN job is available at https://gnps2.org/status?task=37f4e9bbba4e4a65afcdb9496e27d211. Further, MS/MS-containing features were annotated using SIRIUS v6.1 (Dührkop et al., 2019), all possible adducts. The SIRIUS workflow included ZODIAC for molecular-formula identification, CANOPUS for compound-class prediction (Dührkop et al., 2021), NPClassifier for natural-product class assignment (Dührkop et al., 2021), CSI:FingerID (Dührkop et al., 2015) all fall back adducts, structure-similarity searches against the BioCyc, PubChem, HMDB, COCONUT, LOUTS, ChEBI, KEGG databases.

### Statistical analysis of LC-MS/MS

Quantitative feature tables were processed using FBMNstats (Pakkir Shah et al., 2025). Blank subtraction and random-value imputation were applied prior to log₂ transformation. Features were mean-centred across samples. Differential accumulation was tested using the Kruskal–Wallis test followed by Dunn’s post hoc test (p < 0.05).

## Data availability

LC-MS/MS raw and mzXML files for the white flowers (Bride, 72507 and Aristotle) and ResQ flower petals can be downloaded via MASSIVE repository MSV000100849 and MSV000100850 respectively. All the LC-MS/MS data analysis outputs from different tools are provided via zenodo <u>10.5281/zenodo.18641658</u>. RNA-seq data can be obtained from NCBI BioProject ID: PRJNA1426942 with Biosample accessions SAMN55899140-SAMN55899217 and SRA accession numbers are SRR37325350-SRR37325427.

## Results

### Anthocyanin contributes to floral resistance against *B. cinerea*

We quantified total anthocyanin concentrations in the petals from five spontaneous colour mutants of the chrysanthemum cultivar ‘ResQ’. Total anthocyanin concentration followed the visible colour gradient, dark pink and salmon petals had the highest anthocyanin concentrations, while pink and light pink had intermediate amounts, and white petals had no detectable levels of anthocyanins (Fig. 1A). Subsequently, we inoculated the petals with *B. cinerea* and scored for disease incidence, i.e. the number of inoculation droplets that resulted in expanding lesions. Disease incidence showed an inverse relation to anthocyanin concentration: dark pink, salmon, and pink flowers had disease incidence <10%, light pink showed a clear increase to ∼25%, and white flowers were highly susceptible, with disease incidence exceeding 80% (Fig. 1B). This was very clear at 6 dpi, where white petals developed large necrotic lesions, while dark pink petals were completely resistant (Fig. 1C). Together, these data support a negative relationship between petal anthocyanin content and *Botrytis* disease incidence in the *C. morifolium* colour mutants.

**Figure 1:**
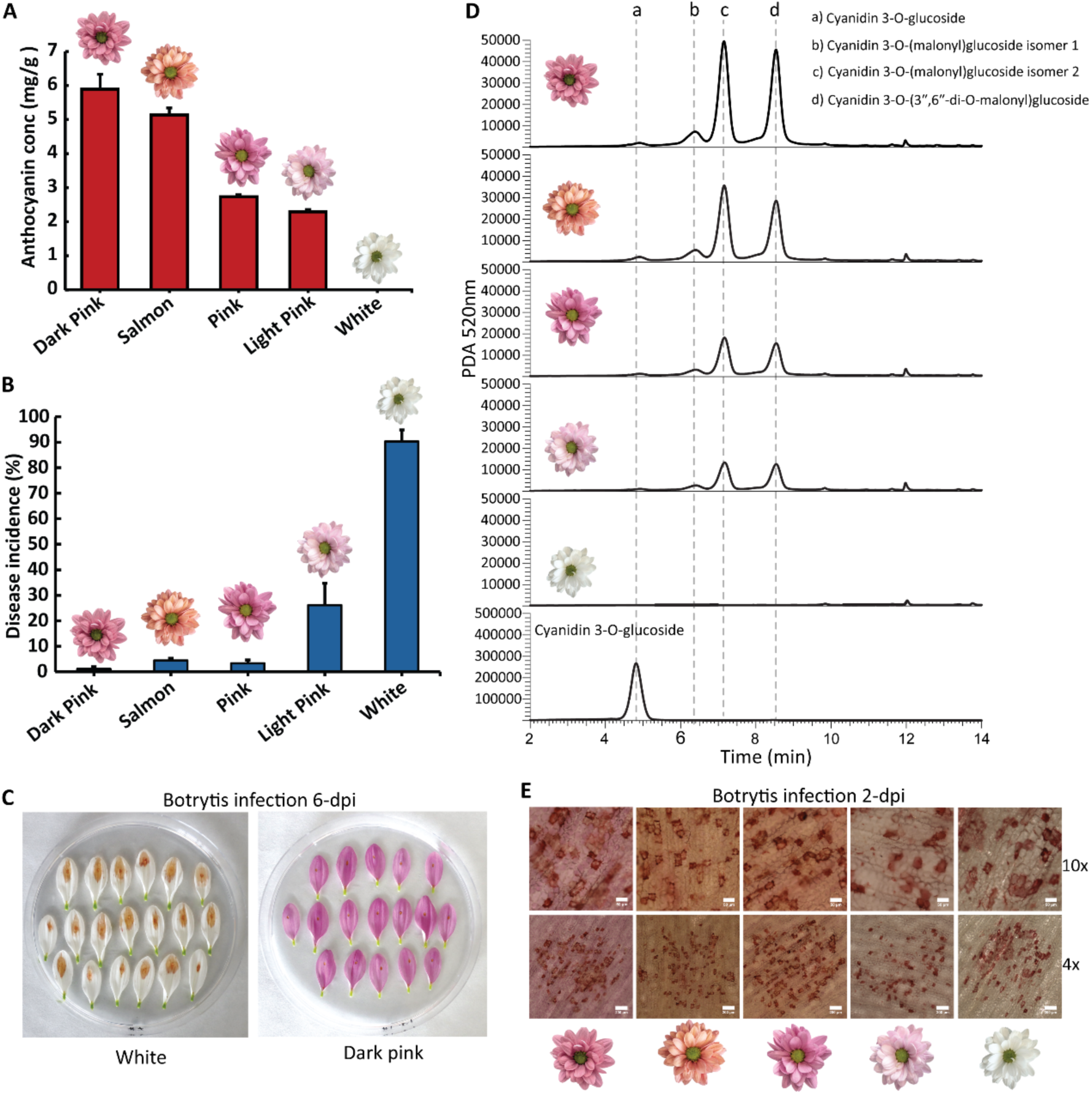
**(A)** Total anthocyanin concentration in petal extracts from dark pink, salmon, pink, light pink, and white ResQ mutants (mean ± SD, n=3). **(B)** Disease incidence after petal inoculation with *B. cinerea* (mean ± SD, n = 180; scoring at 6 dpi). **(C)** Representative petal infection assay at 6 dpi showing extensive necrosis on white petals and no expanding lesions on dark pink petals. **(D)** UHPLC–PDA chromatograms recorded at 520 nm. Vertical dashed lines mark peaks assigned to: **a**, cyanidin 3-O-glucoside; **b**, cyanidin 3-O-(malonyl)glucoside isomer 1; **c**, cyanidin 3-O-(malonyl)glucoside isomer 2; **d**, cyanidin 3-O-(3″,6″-di-O-malonyl)glucoside. Bottom chromatogram is of the cyanidin 3-O-glucoside standard. **(E)** Bright-field microscopy of infected petal tissue at 2 dpi showing red-pigmentation at the inoculation sites. Scale bars: 50 µm (10×) and 200 µm (4×). Flower icons indicate the mutant line.

UHPLC–PDA profiles recorded at 520 nm showed that the coloured lines share the same main anthocyanin peaks, matching with the standard cyanidin 3-O-glucoside and three malonylated derivatives (two mono-malonyl isomers and one di-malonyl form) (Fig. 1D).

Peak heights followed the total anthocyanin concentration. At 2 days post inoculation (dpi), brightfield microscopy revealed discrete red pigmentation at the inoculation sites (Fig. 1E), consistent with the infection-induced, localized anthocyanin accumulation reported for the wild chrysanthemum (*C. seticuspe*) (Suraj et al., 2025). Notably, these red pigments were also observed in *B. cinerea* inoculated white flowers, indicating that infection can trigger red pigment formation even in otherwise non-pigmented petals. Taken together, the data support a strong positive relationship between petal anthocyanin concentration and resistance to *B. cinerea* in the *C. morifolium* petals.

### White chrysanthemum retains infection-triggered red pigmentation response with distinct spatio-temporal patterns

Further we tested multiple diverse white flowered *Chrysanthemum morifolium* cultivars to determine whether they can mount the infection-induced red pigmentation response (Supplementary Figure 1A). All the tested white cultivars produced red pigmentation at the pathogen inoculation sites, and so far, we have not identified a white cultivar that failed to develop this response. However, the magnitude and spatial pattern of pigmentation, as well as disease incidence, varied between cultivars. Three cultivars showed particularly distinct phenotypes (Fig. 2A). ‘Aristotle’ behaved as a hyper-accumulator, producing visible red pigmentation as early as 12 hours post inoculation (hpi) that intensified over-time into dark red patches around inoculation sites over time. In contrast, ‘Bride’ was a slow-accumulator, with delayed and weaker pigmentation that became apparent only from ∼16 hpi. The cultivar ‘72507’ displayed a striking inter-cellular distribution, with pigmentation frequently outlining cell boundaries rather than filling cells, although both intercellular and intracellular pigmentation could be observed in all the three cultivars (Fig. 2A). All three cultivars were resistant to *B. cinerea* under the inoculated conditions used, with disease incidence of 0%.

**Figure 2:**
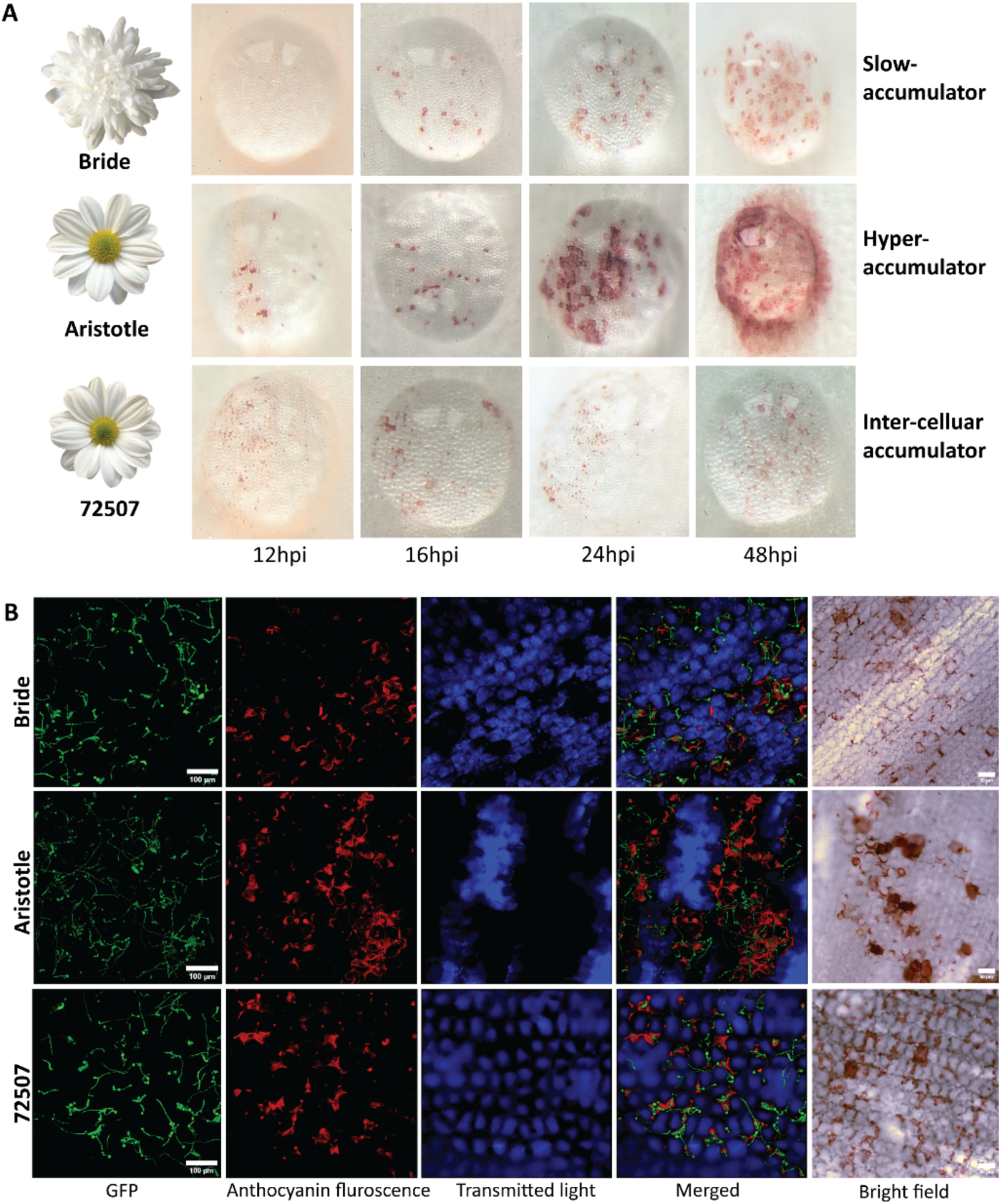
Infection-triggered red pigmentation in three white chrysanthemum cultivars shows cultivar-specific patterns. **(A)** Time course of visible red pigmentation after *B. cinerea* inoculation (12, 16, 20, 48 hpi) in white cultivars ‘Bride’, ‘Aristotle’, and ‘72507’. Cultivars are grouped by the observed response class: slow-accumulator (‘Bride’), slow-accumulator (‘Aristotle’), and inter-cellular accumulator (‘72507’) as seen under stereo-microscope. **(B)** Confocal images from infected petal tissue showing GFP-labelled *B. cinerea* (green), anthocyanin autofluorescence (red), transmitted light (blue), merged channels, and a representative bright-field image for each cultivar at 24 hp (i.e., 1 dpi).

We inoculated the three white cultivars with a GFP-expressing *B. cinerea* strain, and performed confocal microscopy at 24 hpi. Anthocyanin autofluorescence accumulated locally at and around fungal penetration sites (Fig. 2B), consistent with a highly localized host response. Importantly, the cellular context of pigmentation differed: inter-cellular pigmentation was typically observed in tissue where neighbouring cells appeared to retain their structural integrity, whereas intra-cellular pigmentation was often associated with loss of cellular integrity, visible as dark, empty regions in the transmitted-light channel (Fig. 2B). Together, these results show that white flowered cultivated chrysanthemum retains an infection-triggered pigmentation response, but these cultivars differ substantially in both the timing and cellular localization of pigment accumulation.

### Infection activates the flavone and anthocyanin biosynthetic pathway

To connect the visible red pigmentation to an underlying biosynthetic response, we performed transcriptomic and metabolomic analyses. We profiled transcript levels of core flavonoid/anthocyanin pathway genes in three white chrysanthemum cultivars (72507, Aristotle, Bride) at three time points and one time point for the red cultivar (used as a control) after *B. cinerea* inoculations via RNA-seq. Infection caused strong transcriptional activation of the general phenylpropanoid pathway, most clearly seen for the early biosynthetic genes *CHS*, *CHI*, *F3H*, and *F3′H* in the white cultivars, with the largest shifts at later time points (Fig. 3). In contrast, transcript levels of *MYB4*, a negative regulator of the anthocyanin pathway (Zhou et al., 2022) and *FLS* were lower in infected samples than in mock samples (Fig. 3), pointing to reduced flux into the flavonol branch during infection. Transcripts for anthocyanin biosynthetic genes (*DFR*, *ANS, arGST,* and *UFGT*) and *MYB6*, a positive regulator of the anthocyanin pathway in chrysanthemum (Xia et al., 2026) increased in the white cultivars after *B. cinerea* inoculations (Fig. 3; Supplementary Figure 2B). This pattern was consistent with the spatio-temporal nature of the red pigmentation response, suggesting that anthocyanin pathway activation is localized to fungal penetration sites rather than broadly induced across the petal tissue. As expected, the red-flowered control showed stronger expression of late-pathway genes than the white cultivars (Fig. 3), matching its stronger pigment capacity.

**Figure 3:**
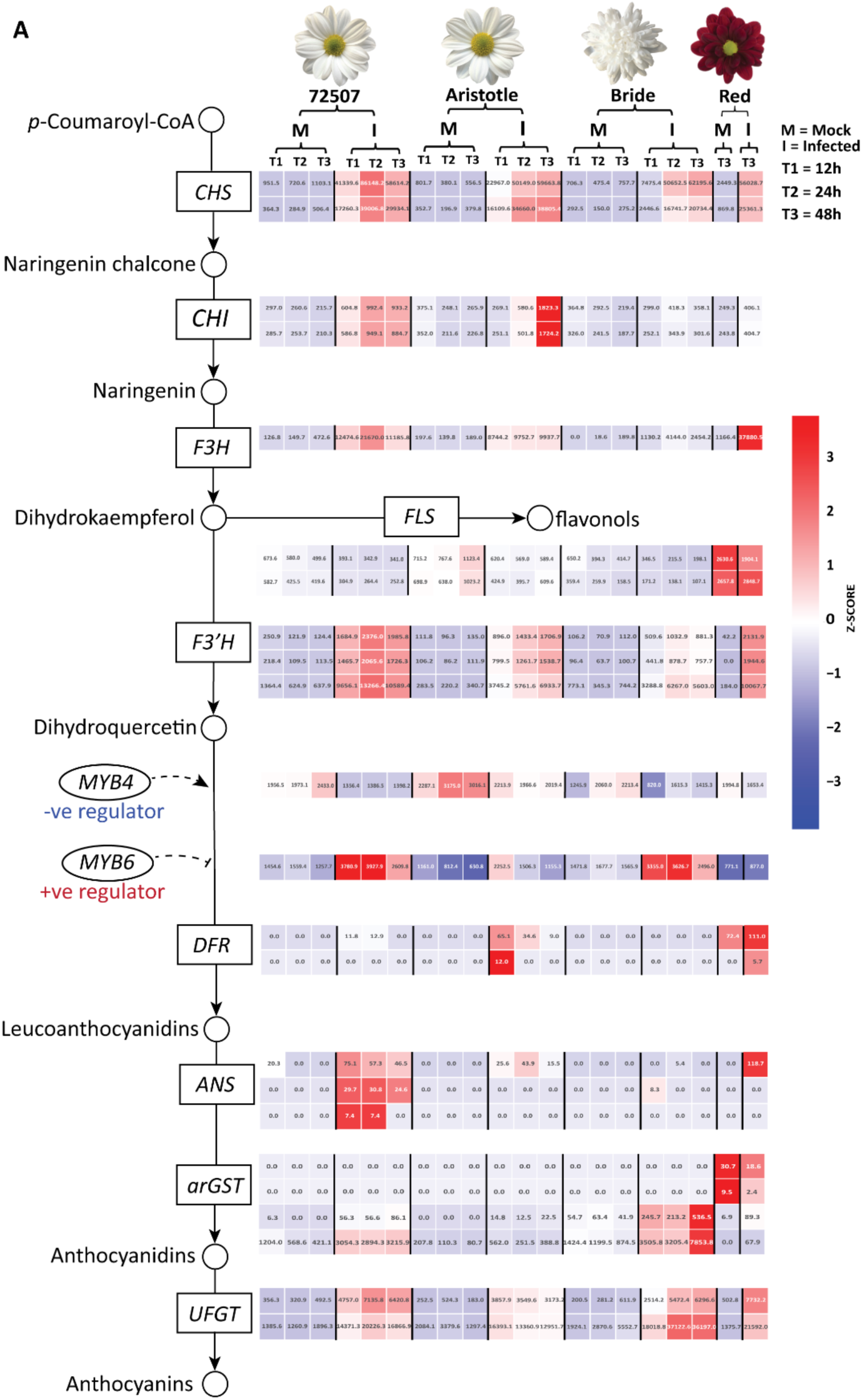
Heatmap of flavonoid/anthocyanin biosynthetic gene expression in Chrysanthemum petals after *B. cinerea* infection. Transcript levels of flavonoid/anthocyanin pathway genes in three white-flowered *Chrysanthemum* cultivars (72507, Aristotle, Bride) sampled after mock treatment (M) or *B. cinerea* inoculated (I) conditions at 12 h (T1), 24 h (T2), and 48 h (T3) post inoculation. A red cultivar (Mexicano) was included as a control (mock and infected at 24 h). Genes shown: *CHS*, chalcone synthase; *CHI*, chalcone isomerase; *F3H*, flavanone 3-hydroxylase; *F3′H*, flavonoid 3′-hydroxylase; *DFR*, dihydroflavonol 4-reductase; *ANS*, anthocyanidin synthase; *arGST,* anthocyanin-associated glutathione S-transferase; *UFGT*, UDP-glucose flavonoid 3-O-glucosyltransferase; *FLS*, flavonol synthase; MYB4 and MYB6 are R2R3-MYB transcription factor. Cell colours represent row-wise z-scores (blue to red indicates low to high relative expression for each gene). Numbers inside cells are DESeq2-normalized expression values.

In parallel with this, multiple flavone synthase II (*FNS2)* candidate genes were upregulated after infection in all three white cultivars (Supplementary Figure 2B), indicating of activation of the flavone branch. In the same samples, several candidate ABCC transporters were upregulated (Supplementary Figure 2B), indicating that transport/sequestration capacity increases along with the pathway activity. DMACA staining was performed to detect proanthocyanidins, but there was no positive staining for these metabolites in the infected petals. Furthermore, the transcript of *MYB123*, a positive regulator of proanthocyanidin (PA) biosynthesis was undetectable (Supplementary Figure 2C), indicating that PA biosynthesis might not be active in the tested conditions.

Untargeted LC–MS/MS and feature-based molecular networking (FBMN) showed that infection shifts the petal metabolome in a largely similar way, while still displaying cultivar-specific differential metabolite features. In total, 649 metabolite features changed in abundance after infection in at least one cultivar, with 393 features shared by all the three cultivars (Aristotle, 72507, and Bride) (Fig. 4A). Each cultivar also showed a relatively small number of unique features (Fig. 4A). Chemical compound class assignments by SIRIUS-CANOPUS placed most differentially abundant features in the shikimates and phenylpropanoids (Fig. 4B). FBMN analysis showed clear differential abundance in the infected petals for identified anthocyanins, mainly cyanidin-based glycosides and acylated derivatives. Flavones such as apigenin, apigenin-7-glucoside, and tilianin also increased in abundance (Fig. 4C–F). Altogether, the transcript and metabolite data supports a model in which botrytis infection suppresses the flavonol branch, and at the same time activates flavone branch and anthocyanin biosynthesis particularly at the pathogen penetration site.

**Figure 4:**
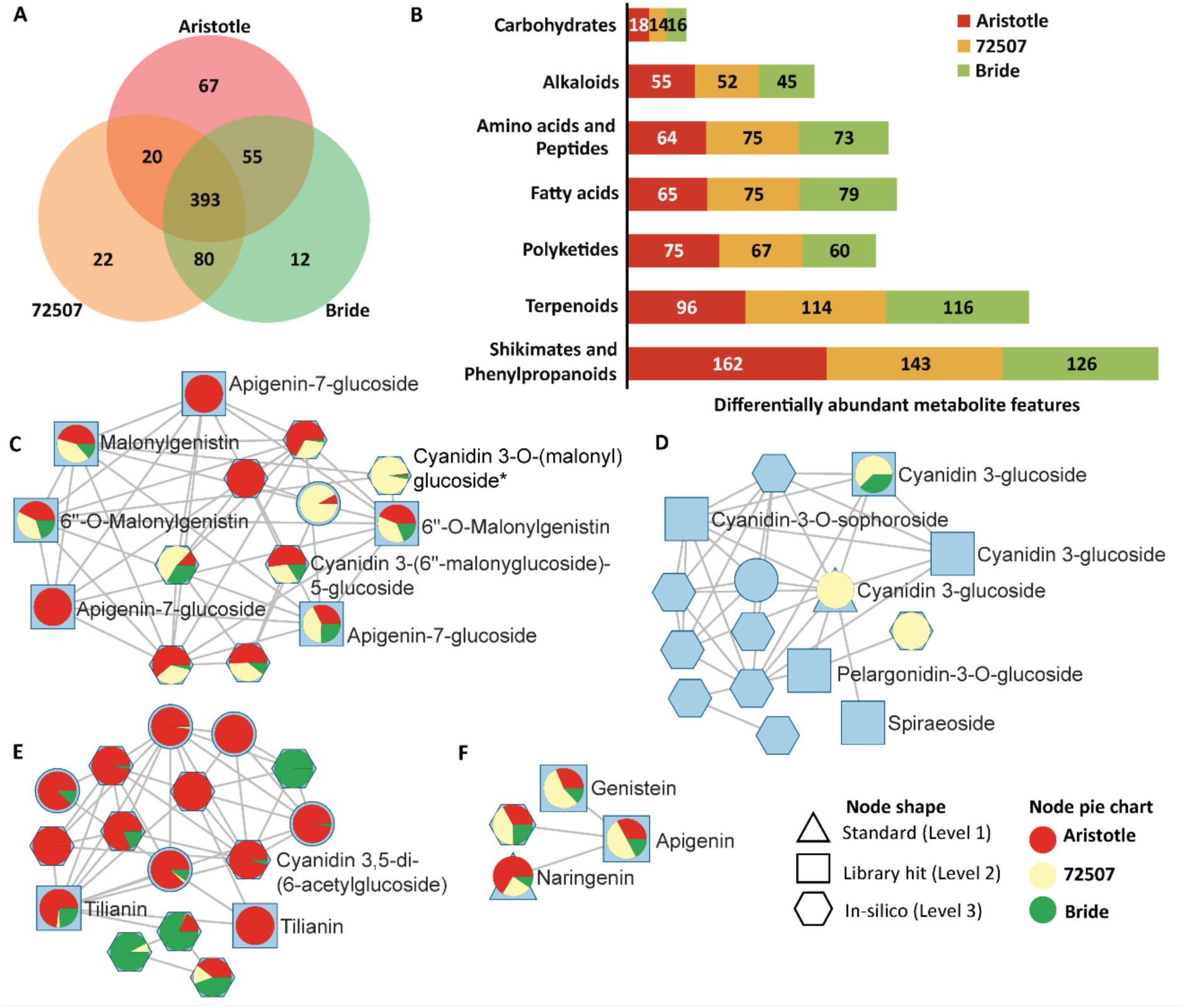
Metabolite changes in three white chrysanthemum cultivars during infection by *B. cinerea*. **(A)** Venn diagram showing overlap in differentially abundant metabolite features between the three white cultivars (Aristotle, 72507, Bride) after *B. cinerea* infection; numbers indicate feature counts per set and overlap. **(B)** Compound class distribution of differentially abundant features annotated by SIRIUS and grouped by NPClassifier chemical superclass. Numbers inside bars indicate feature counts per cultivar (Aristotle, 72507, Bride). **(C–G)** Selected feature-based molecular networking (FBMN) subnetworks from major flavonoid/anthocyanin-related molecular families. Nodes represent individual LC–MS features (defined by precursor *m/z* and retention time), and edges represent MS/MS spectral similarity. All displayed nodes correspond to features that were significantly more abundant in infected samples. Node pie charts show relative abundance by cultivar (Aristotle = red, 72507 = yellow, Bride = green). Node shape indicates annotation confidence: triangle = authentic standard (Level 1), square = spectral library match (Level 2), hexagon = *in silico* annotation (Level 3); * indicates manual curation. The same metabolite annotation can appear on multiple nodes because FBMN nodes correspond to features, not unique chemical structures. Thus, one compound may be represented by multiple detected forms (e.g., different adducts/ion species, in-source fragments, dimers) and/or by chromatographically resolved isomers that share similar MS/MS-annotation. Only representative labels are shown in the figure for readability. The complete node-level annotation list is provided in an annotation table and the full network files are available via GNPS job link for both see data availability section.

### Infection-induced red pigmentation is restricted to Asteraceae flowers in a cross-angiosperm survey

To test whether infection-triggered red pigmentation is a general property of white flowers or a lineage-specific response, we expanded the petal infection assay to a broad panel of white or pale-flowered angiosperms. Upon inoculation with *B. cinerea*, we monitored for the occurrence of localized red pigmentation versus the browning/tissue clearing that is commonly associated with necrosis (Fig. 5; Supplementary Figure 3). Across the taxa tested, red pigmentation was observed only in Asteraceae flowers with the exception of *Gerbera x hybrida*, *for which* both coloured and white flowers turned brown (Supplementary Figure 4). Red pigments formed in *Chrysanthemum morifolium*, *Leucanthemum vulgare* (wild daisy), *Cosmos bipinnatus*, and *Helianthus annuus* (wild sunflower) after *B. cinerea* infection (Fig. 5A,C). In contrast, all non-Asteraceae species examined did not develop red pigmentation and instead showed browning/tissue clearing at the infection sites (Fig. 5C; Supplementary Figure 3). Mapping these outcomes onto a simplified phylogeny highlighted a clear phylogenetic signature, consistent with an Asteraceae-associated capacity for infection-triggered red pigment formation in otherwise non-pigmented or yellow petals (Fig. 5C). Notably, pigment induction aligned with disease outcome, the wild daisy and wild sunflower produced red pigment after infection and showed lower disease incidence whereas the cultivated species did not show this response and were highly susceptible to *B. cinerea*.

**Figure 5:**
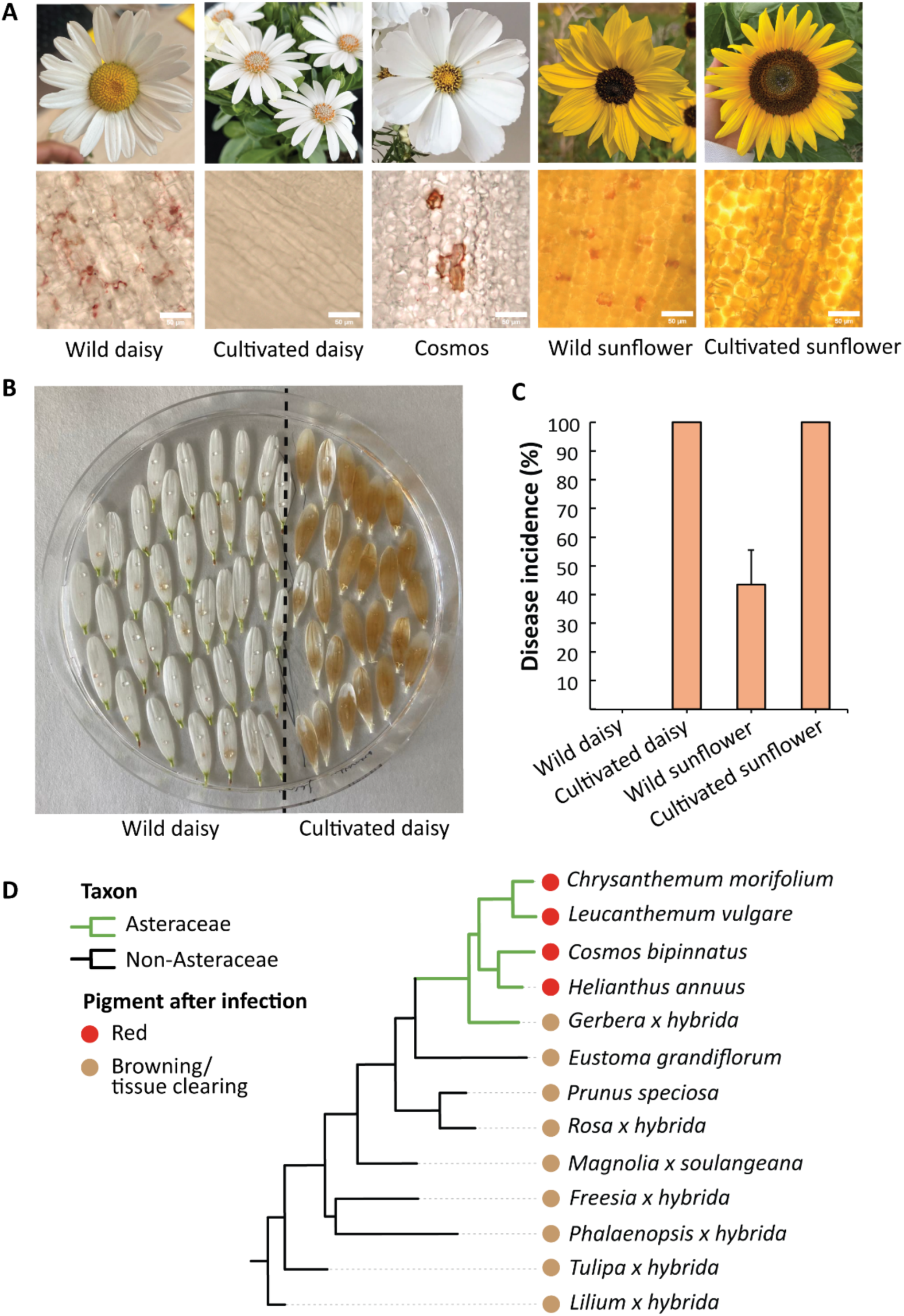
Infection-induced red-pigment response in white flowers are common in several Asteraceae species but rare outside the family. **(A)** Botrytis infection on wild daisy, cultivated daisy, cosmos, wild sunflower and cultivated sunflower (1-dpi). Scale bars, 50 µm. **(B)** Representative detached-petal assay | 3-dpi after infection. The wild daisy is resistant to *B. cinerea* and shows red-pigmentation response while the opposite observed in cultivated daisy. **(C)** Disease incidence of wild vs cultivated daisy/sunflower after inoculations with *B. cinerea* 3-dpi, the error bars represents standard deviation with n = 40. **(D)** Schematic phylogeny of tested taxa. Branch colour indicates taxonomic group (Asteraceae vs non-Asteraceae). Symbols indicate the dominant visible outcome after infection: red pigment (red circles) or browning/tissue clearing (brown circles).

## Discussion

This study utilized a unique genetic resource consisting of spontaneous colour mutants (ResQ lines) derived from a single breeding lineage. Since these lines are near-isogenic and differ primarily in petal anthocyanin concentration, they provide a unique opportunity to evaluate the direct impact of pigmentation on susceptibility of flowers to *B. cinerea* with minimal background genetic noise. Consistent with the previous reports that anthocyanins contribute to plant defense against both biotic and abiotic stress (Grünig et al., 2025), we observed that highly pigmented flowers exhibited increased resistance, whereas the white flowers were highly susceptible to *B. cinerea*. The colour difference within the ResQ panel is likely driven by mutations in the regulators of the anthocyanin pathway, with MYB-family transcription factors serving as primary candidates. Our findings align with previous reports suggesting a high abundance of flavonols/flavones in white chrysanthemum flowers (Chen et al., 2012). Here, the white ResQ flowers showed significantly higher *FLS* and *FNS2* expression compared to differentially coloured lines (Supplementary Figure 2A). However, the expression patterns of anthocyanin biosynthetic genes presented an intriguing paradox. While one might expect high *DFR* and *ANS* expression in coloured petals, expression of these genes was nearly undetectable in many of the coloured ResQ flowers. Even when screened across multiple developmental stages (early buds to mature flowers) *ANS* gene expression measured via qPCR remained undetectable (Supplementary Figure 6). However, we observed that early buds were already pigmented, suggesting that anthocyanin biosynthesis in these flowers are likely governed by a negative feedback loop mechanism (LaFountain & Yuan, 2021; Yan et al., 2021). This is further supported by the elevated expression of the negative regulator *MYB4* in the petals of coloured ResQ lines (Supplementary figure 2A), which implies lack of activation of anthocyanin biosynthesis once a specific pigment threshold is reached.

A particularly striking result of this study was the localized induction of red pigmentation in petals of multiple white cultivars following *B. cinerea* inoculations. This pattern suggests the occurrence of a site-specific stress response at pathogen penetration sites. The metabolomic data confirmed that these red pigments consist of *de novo* synthesized anthocyanins. Remarkably, pigmentation was observed in both intra-cellular and inter-cellular (apoplastic) spaces. Since anthocyanins are typically considered to be unstable outside the acidic environment of the vacuole (Alcalde-Eon et al., 2024), their presence in the apoplast suggests a stabilization mechanism. We hypothesize that this stability is mediated by two, potentially synergistic factors: co-pigmentation and pathogen-induced acidification. First, the white chrysanthemum flowers maintain high basal levels of flavones (Chen et al., 2012), including apigenin 7-O-glucoside, which are further bolstered by the pathogen-induced upregulation of flavones (Supplementary figure 2B). These flavones can form non-covalent complexes with anthocyanins (via stacking or sandwich model), stabilize anthocyanins and protect them against nucleophilic water attack (Dangles et al., 1993; Alcalde-Eon et al., 2024). Co-pigmentations mechanism is well studied in red wine and it is responsible for 30–50% of the coloration in young red wines (Trouillas et al., 2016). This mechanism is further supported by studies in the blue chrysanthemums, it was found that blue colouration depends on the co-pigmentation of anthocyanins with flavones (Noda et al., 2017). Simultaneously, early *B. cinerea* infection is typically associated oxalic acid secretion and lowering of local apoplastic pH (Williamson et al., 2007). Acidic pH help stabilize the red flavylium cations and thereby contribute to retaining a visible red colour (Kallam et al., 2017). Future research should focus on understanding such mechanisms in detail as it provides opportunity for making designer flowers, a step forward in flower engineering.

The evolutionary transition from pigmented to non-pigmented flowers is a long-studied question in plant biology. Current evidence suggests that the loss of flower pigmentation is frequently driven by regulatory changes affecting MYB transcription factors (Marin-Recinos & Pucker, 2024; Wheeler et al., 2022). This regulation is further dependent on the substrate competition, particularly between *FLS* and *DFR.* The substrate competition is mitigated through conserved amino acid residues (His-132 in *FLS* and Asp-133 in *DFR*), which bias the substrate preference (Choudhary & Pucker, 2024; Johnson et al., 2001). To prevent metabolic conflict, plants exhibits reciprocal expression patterns, where the anthocyanin producing *DFR* is upregulated only when needed, while *FLS* dominates otherwise (Choudhary & Pucker, 2024). This allows the plant to flexibly direct metabolic flux towards either flavonols or anthocyanins depending on the developmental or environmental cues. Our findings in the white chrysanthemum flowers align with this model of competitive regulation. Though these flowers appear white and chemically favor flavone/flavonol production under normal conditions, the structural genes for anthocyanin biosynthesis remain intact. Upon *B. cinerea* infection, we observe an inversion of this regulation (downregulation of *FLS* and upregulation of *DFR*), triggering anthocyanin biosynthesis specifically at the pathogen penetration sites. Thus, the white phenotype in chrysanthemum does not represent a loss of genetic potential, but rather a dormant state that can be rapidly overridden by environmental stress cues.

The phenomenon of pathogen-induced pigmentation is not unique to chrysanthemum. Similar localized anthocyanin accumulation has been documented in other non-pigmented tissues, such as white grapes, white corolla tubes of *Antirrhinum* and white rose petals following infections with *B. cinerea* (Blanco-Ulate et al., 2015; Harrison & Stickland, 1980; Li et al., 2025). These observations suggest that anthocyanins, with their well-documented antioxidant and antimicrobial properties (Deng et al., 2025), function as a basal defence and delays cell death to limit pathogen expansion. To determine if such response is universal or pathogen specific, we inoculated white flower petals of chrysanthemum with pathogens exhibiting diverse lifestyles: the biotroph *Fulvia fulva*, the hemi-biotroph *Fusarium oxysporum* f.sp. *chrysanthemi*, and the necrotroph *Alternaria solani* (Supplementary figure 6). Interestingly, only *A. solani* triggered a red pigmentation response similar to the one observed upon *B. cinerea* inoculations. This indicates that the induction of the anthocyanin pathway in chrysanthemum is not a generalized stress response, but might rather be a specific response to necrotrophic pathogens. We speculate that the plant recognizes specific effectors common to this class of pathogens that trigger the anthocyanin pathway. While preliminary tests with a filtered extract from the *B. cinerea* infection site in *Antirrhinum* indicated the presence of a low-molecular weight effector (Harrison & Stickland, 1980), stronger evidence for the existence/identification of this pigment-inducing factor requires follow-up studies.

To assess the evolutionary conservation of this trait, we surveyed across a broader range of non-pigmented flowers. While this response is only observed sporadically in other branches of flowering plants (Harrison & Stickland, 1980; Li et al., 2025), the mechanism seems conserved within the Asteraceae family with the notable exception of gerbera, regardless of their colour (Supplementary figure 3 and 4). This could be the result of loss-of-function in *Gerbera* or a gain-of-function in the Asteraceae clade after *Gerbera* diverged. The red pigmentation response was absent in other plant families tested including the Solanaceae. Infection of multiple *Petunia* genotypes (both wild-type and flavonoid mutants) failed to elicit red pigmentation (Supplementary figure 5), suggesting that the regulators of anthocyanin biosynthesis in *Petunia* does not respond to infection from the necrotrophic pathogen. Our results revealed a potential domestication trade-off within the Asteraceae. While wild species of Asteraceae frequently produced red pigmentation and showed higher resistance, many cultivated species such as commercial daisies and sunflowers lacked this response and were significantly more susceptible to *B. cinerea*. This suggests that breeding for aesthetic traits (for example, pure white flowers) may have inadvertently selected against this stress-responsive regulatory module. Future work should determine what factors in the pathogen triggers this response, and unravel the regulatory mechanisms controlling this response, as well as pinpoint how this mechanism was lost during breeding. This could help in breeding white flowers that can respond to infection and produce anthocyanins to stop the spread of infections.

## Acknowledgements

We thank all members of the Phytopathology Laboratory of WUR for technical assistance and helpful discussions. We acknowledge the support from Dr. Ric de Vos (Wageningen University & Research, The Netherlands) in the metabolomics work. We are grateful to Dr. Nick Albert (Plant & Food Research, New Zealand) and Dr. Cathie Martin (John Innes Centre, UK) for their valuable scientific input and Dr. Francesca Quattrocchio (University of Amsterdam, The Netherlands) for providing petunia mutant flower collection. Felicia Wolters (Wageningen University & Research, The Netherlands) for assistance and support in analysing metabolomics data. HS acknowledges funding from the TKI Graduate School Green Top Sectors, The Netherlands (grant LWV21.291, TU-2021-20). We thank Aike Post (Deliflor Chrysanten) and Dr. Nick de Vetten (Dekker Chrysanten) for fruitful discussions during project meetings, and we acknowledge Deliflor Chrysanten and Dekker Chrysanten for the supply of plant materials and for co-funding the project together with the TKI grant.

## Author contributions

HS contributed to conceptualisation, investigation, data analysis, visualisation, and writing—original draft, review, and editing. LR contributed to investigation, data analysis, visualisation, and writing—review and editing. JJJvdH contributed to methodology, supervision, manuscript drafting, and writing—review and editing. SM contributed to conceptualisation, visualisation, and writing—review and editing. BP contributed methodology, data analysis, visualisation and writing—review and editing. JK contributed to conceptualisation, methodology, visualisation, project administration, supervision, and writing—review and editing.

## Competing interests

JJJvdH is member of the Scientific Advisory Board of NAICONS Srl., Milano, Italy and consults for Corteva Agriscience, Indianapolis, IN, USA. All other authors declare to have no competing interests.

## Supplementary figures

**Supplementary Figure 1:**
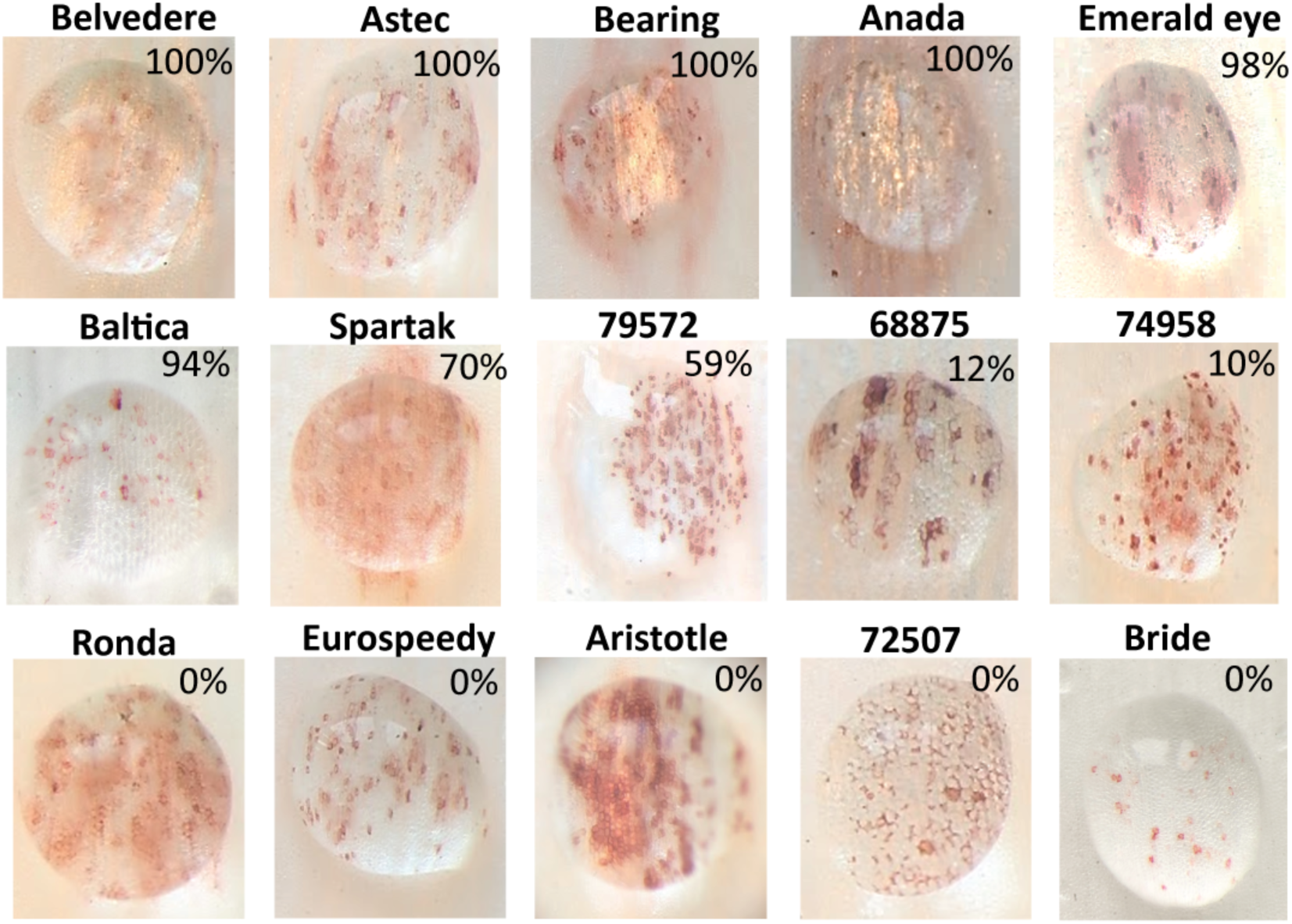
Infection-induced red pigmentation occurs across diverse white chrysanthemum cultivars. Representative infected petal images from a panel of white-flowered *Chrysanthemum morifolium* cultivars showing variable formation of localized red pigmentation 24 hrs after *B. cinerea* inoculation. Percentages indicate disease incidence for each cultivar, scored as the number of expanding lesions 6 days post inoculations: 100% denotes complete susceptibility (all petals diseased), whereas 0% denotes no expanding lesions (fully resistant) n=30.

**Supplementary Figure 2:**
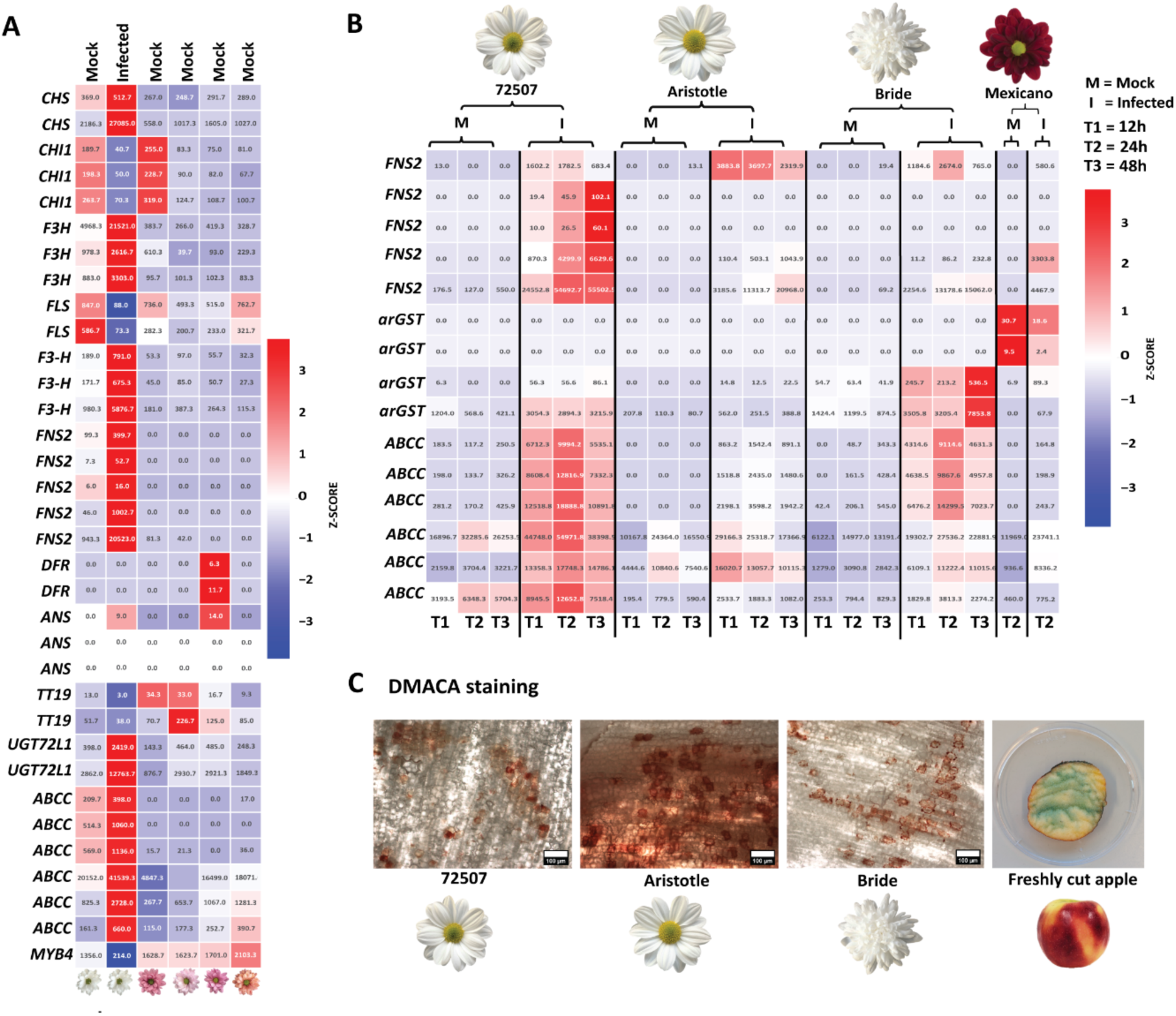
**(A)** Heatmap of expression measured via RNA-seq for flavonoid/anthocyanin genes in the ResQ colour series (mock vs infected (2-dpi, white petals), shown as row-wise z-scores with DESeq2-normalized expression values overlaid in each cell. **(B)** Heatmap of RNA-seq expression in three white cultivars (72507, Aristotle, Bride) and a red cultivar across a 12–48 h time course after *B. cinerea* inoculation (mock, M; infected, I). Genes shown include *FNS2*, *arGST*, and multiple *ABCC* transporter candidates, displayed as row-wise z-scores with DESeq2-normalized expression values overlaid. Time points: T1 = 12 h, T2 = 24 h, T3 = 48 h post inoculation. **(C)** DMACA staining of infected petals from the three white cultivars showing no positive proanthocyanidin signal; freshly cut apple tissue is included as a positive staining control.

**Supplementary Figure 3:**
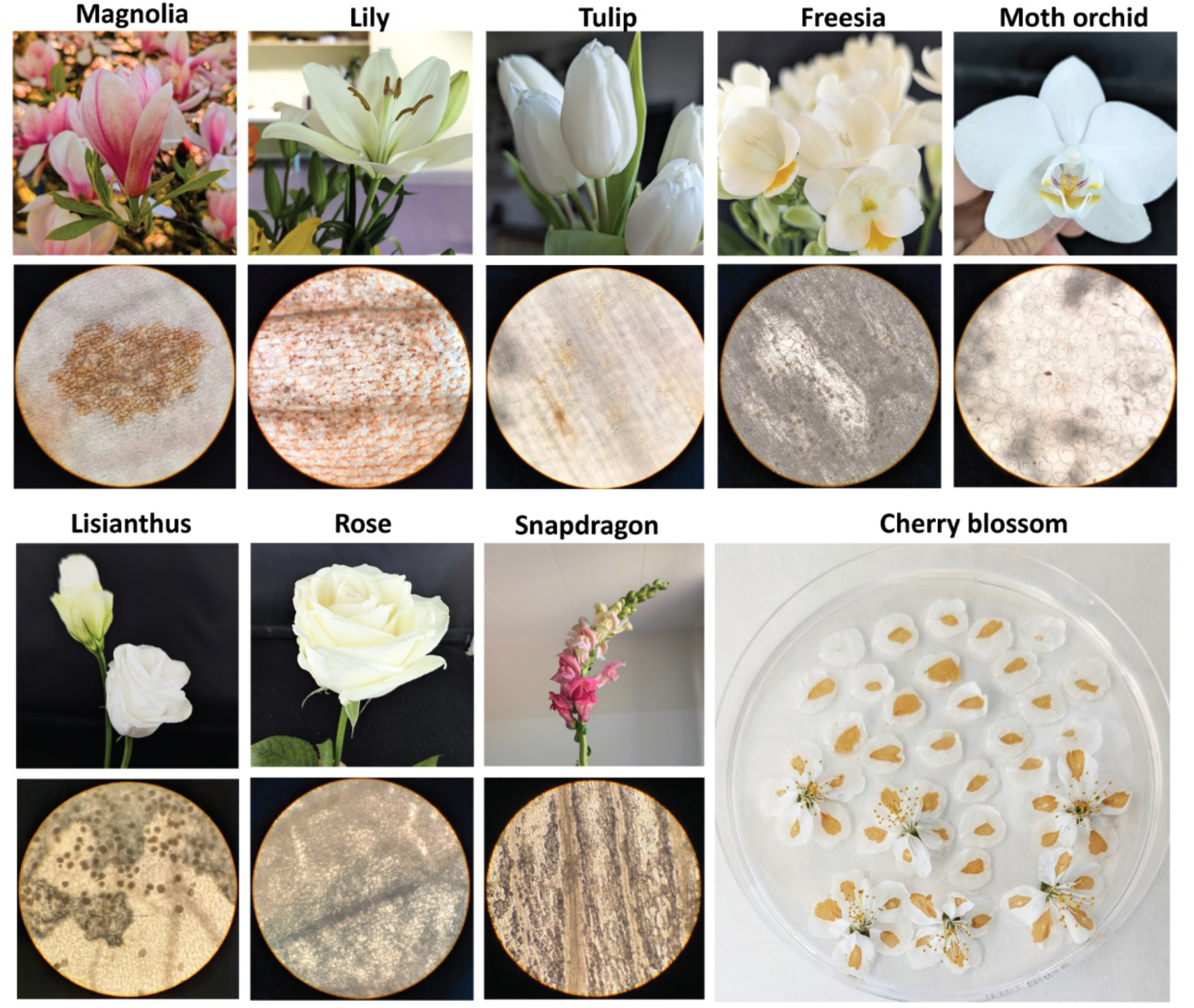
Representative flower images (top) and the stereomicroscope images of the corresponding petals at inoculation sites (bottom) 24 hrs after *Botrytis cinerea* inoculations for white flowers, including Magnolia (*Magnolia soulangeana*), lily (*Lilium × hybrida*), tulip (*Tulipa x hybrida*), freesia (*Freesia × hybrida*), moth orchid (*Phalaenopsis × hybrida*), Snapdragon (*Antirrhinum majus*), lisianthus (*Eustoma grandiflorum*), rose (*Rosa × hybrida*) and the cherry blossom (*Prunus speciosa*) is shown as a detached-flower assay plate with severe necrotic lesions after infection.

**Supplementary Figure 4:**
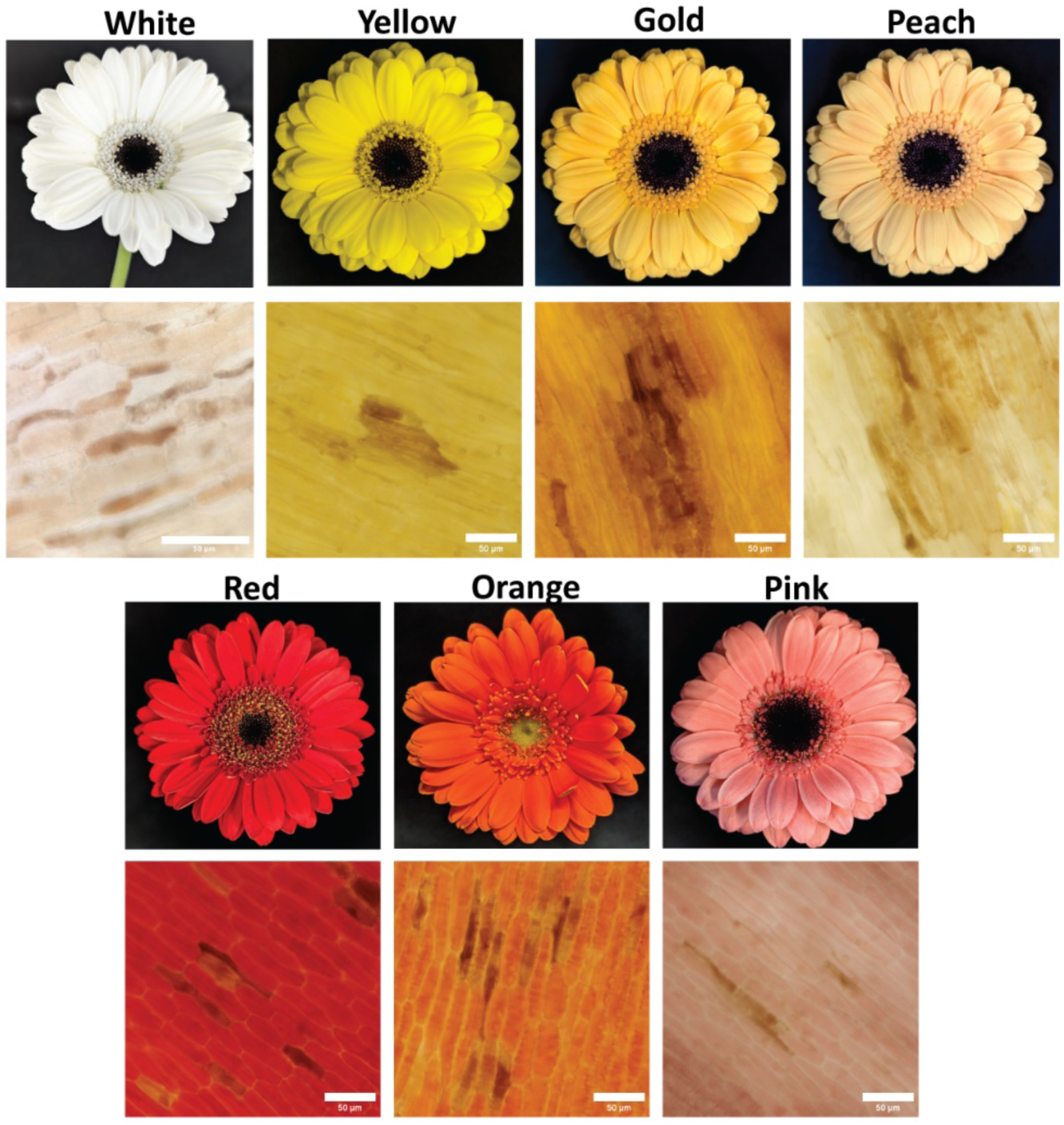
Representative flower images (top) and the stereomicroscope images of the corresponding petals at inoculation sites (bottom) 24 hrs after *B. cinerea* inoculations for white and coloured Gerbera (*Gerbera x hybrida*) showing browning/necrosis.

**Supplementary Figure 5:**
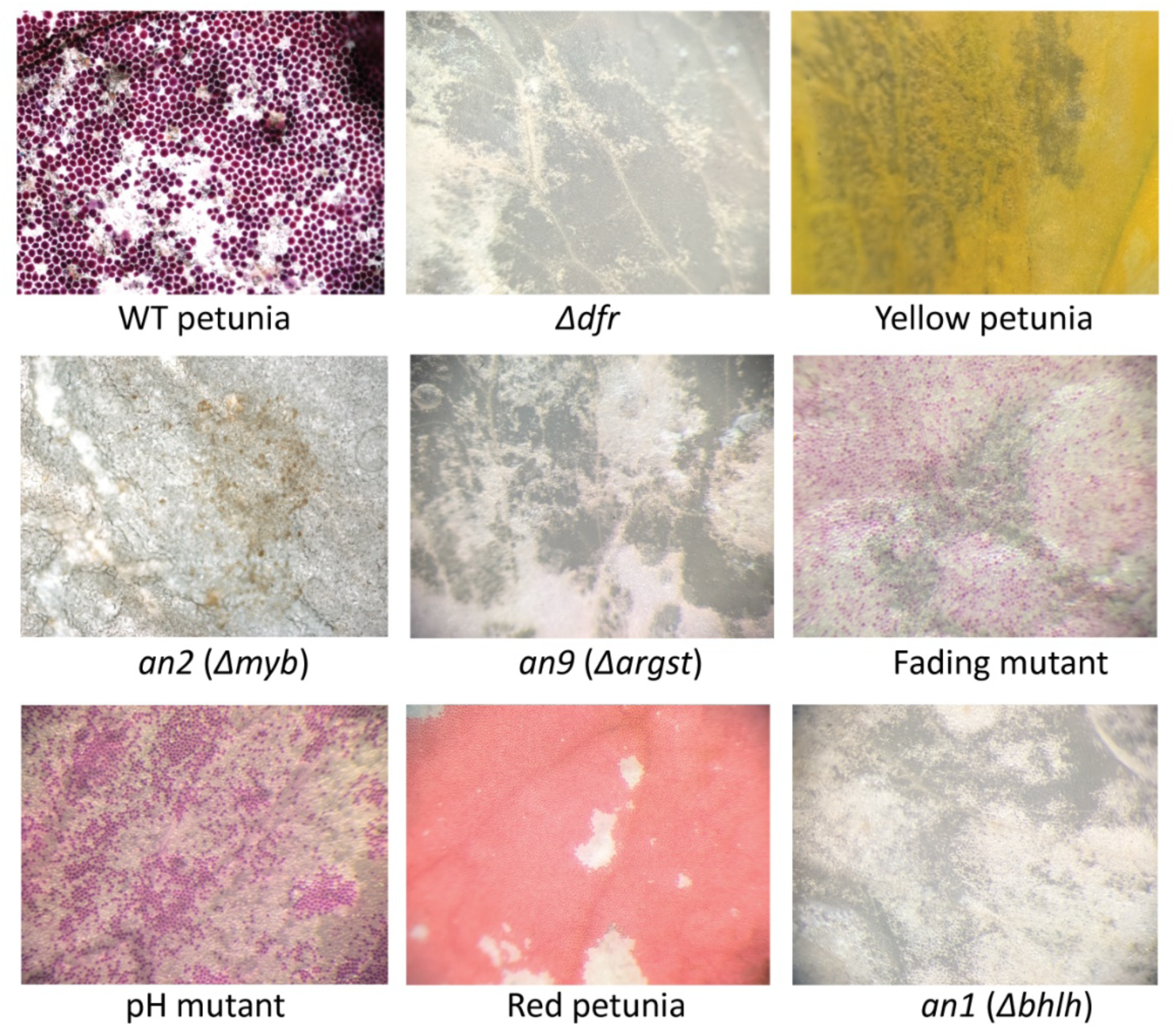
Representative images of inoculated petal regions 24 hpi after *B. cinerea* in wild-type and anthocyanin/flavonoid mutants of petunia. Genotypes include *Δdfr* (dihydroflavonol 4-reductase mutant), yellow petunia, *an2* (*Δmyb*) (R2R3-MYB regulator mutant), *an9* (*Δargst*) (anthocyanin-associated glutathione S-transferase mutant), fading mutant, pH mutant (vacuolar pH-related colour mutant), Red petunia (pigmented control), and an1 (Δ*bhlh*) (bHLH regulator mutant).

**Supplementary Figure 6:**
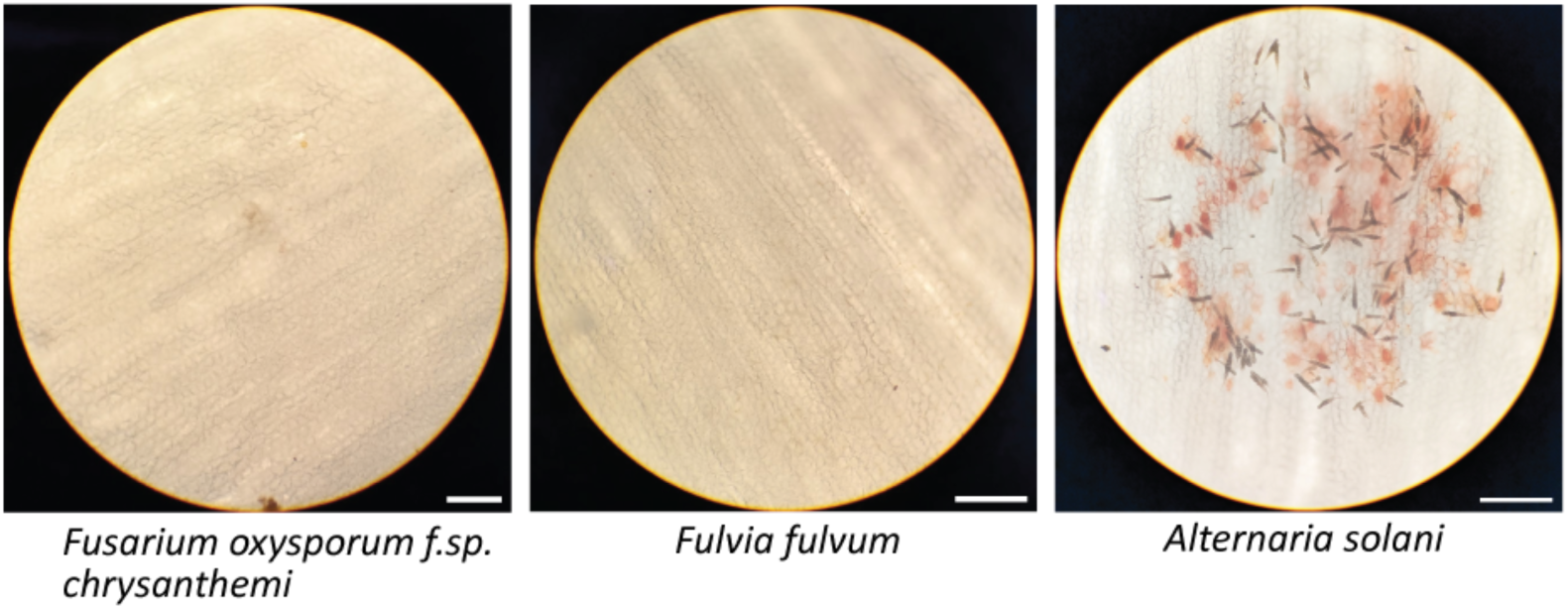
Microscopy of inoculated petal tissue with different fungal pathogens 24 hpi. *Fusarium oxysporum* and *Fulvia fulvum* did not induce red pigmentation, whereas *Alternaria solani* triggered visible red pigmentation at infection sites under the same assay conditions. Scale bars are 200 μm.

**Supplementary Figure 7:**
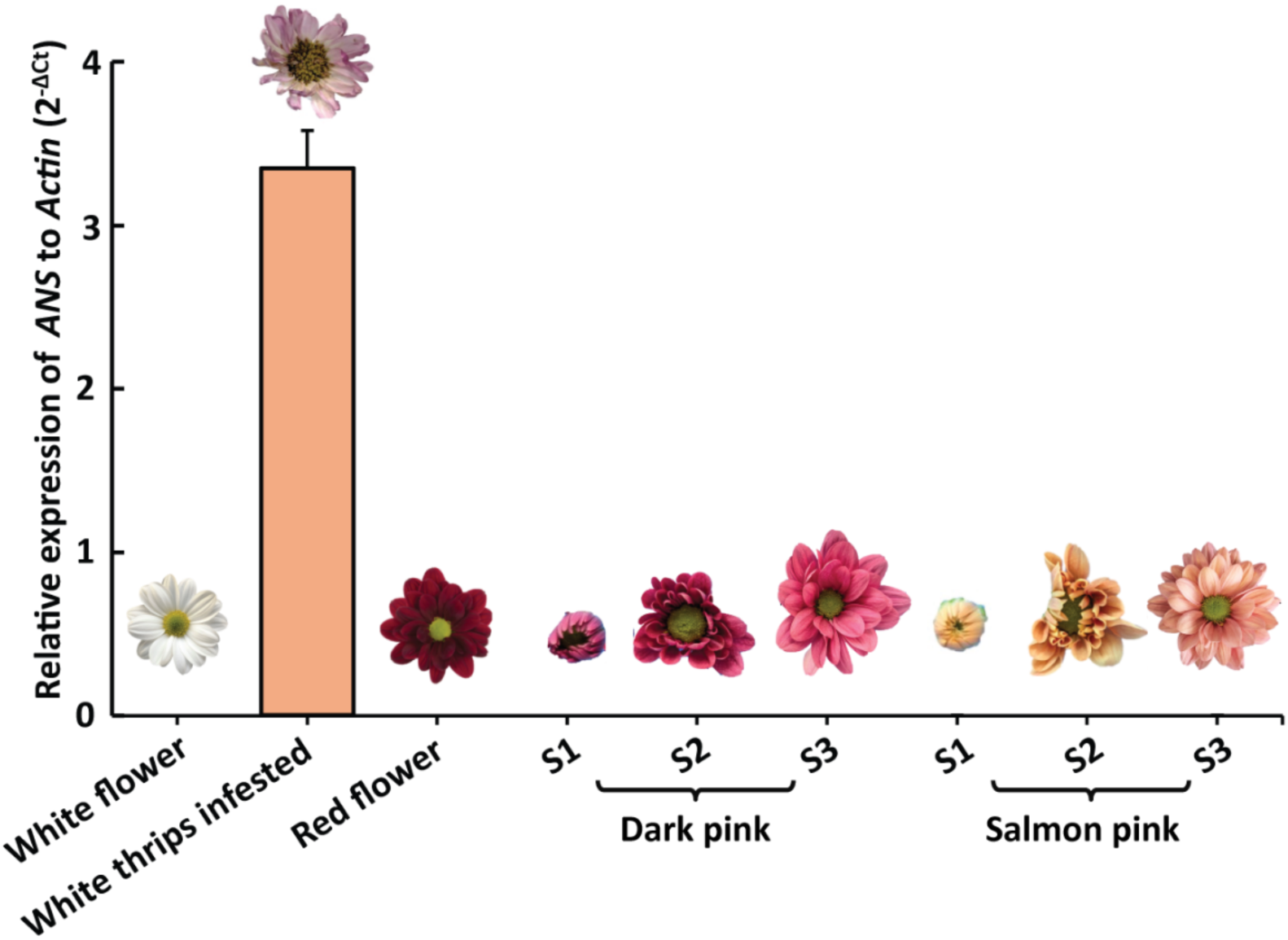
Relative *ANS* transcript levels were quantified by qPCR, normalized to *Actin*. Dark-pink and salmon flowers: S1–S3 represent flower development stages: S1 (closed bud), S2 (partially open flower), and S3 (fully open flower). Mature white and mature red flowers are included for comparison. A thrips-infested white flower with induced anthocyanin pigmentation is used as a positive control. Error bars represent standard deviation, n= 3.

